# Global genomic population structure of *Clostridioides difficile*

**DOI:** 10.1101/727230

**Authors:** Martinique Frentrup, Zhemin Zhou, Matthias Steglich, Jan P. Meier-Kolthoff, Markus Göker, Thomas Riedel, Boyke Bunk, Cathrin Spröer, Jörg Overmann, Marion Blaschitz, Alexander Indra, Lutz von Müller, Thomas A. Kohl, Stefan Niemann, Christian Seyboldt, Frank Klawonn, Nitin Kumar, Trevor D. Lawley, Sergio García-Fernández, Rafael Cantón, Rosa del Campo, Ortrud Zimmermann, Uwe Groß, Mark Achtman, Ulrich Nübel

## Abstract

*Clostridioides difficile* is the primary infectious cause of antibiotic-associated diarrhea. Local transmissions and international outbreaks of this pathogen have been previously elucidated by bacterial whole-genome sequencing, but comparative genomic analyses at the global scale were hampered by the lack of specific bioinformatic tools. Here we introduce EnteroBase, a publicly accessible database (http://enterobase.warwick.ac.uk) that automatically retrieves and assembles *C. difficile* short-reads from the public domain, and calls alleles for core-genome multilocus sequence typing (cgMLST). We demonstrate that the identification of highly related genomes is 89% consistent between cgMLST and single-nucleotide polymorphisms. EnteroBase currently contains 13,515 quality-controlled genomes which have been assigned to hierarchical sets of single-linkage clusters by cgMLST distances. Hierarchical clustering can be used to identify populations of *C. difficile* at all epidemiological levels, from recent transmission chains through to pandemic and endemic strains, and is largely compatible with prior ribotyping. Hierarchical clustering thus enables comparisons to earlier surveillance data and will facilitate communication among researchers, clinicians and public-health officials who are combatting disease caused by *C. difficile*.

## Introduction

The anaerobic gut bacterium *Clostridioides difficile* (formerly *Clostridium difficile*) ^1^ is the primary cause of antibiotic-associated diarrhea in Europe and North America ^2^. Molecular genotyping of *C. difficile* isolates has demonstrated international dissemination of diverse strains through healthcare systems ^3-5^, the community ^6^, and livestock production facilities ^7,8^. Previously, genotyping was commonly performed by PCR ribotyping or DNA macrorestriction. More recent publications have documented that genome-wide single-nucleotide polymorphisms (SNPs) from whole-genome sequences provide improved discrimination, and such analyses have enabled dramatic progress in our understanding of the emergence and spread of pandemic strains ^9-12^ and the epidemiology of local transmission ^13,14^. Eyre and colleagues have argued that transmission of *C. difficile* isolates within a hospital environment can be recognised with high probability as chains of genomes which differ by up to two SNPs whereas genomes which differ by at least 10 genomic SNPs represent unrelated bacteria ^13,15^. However, SNP analyses require sophisticated bioinformatic tools and are difficult to standardize ^16,17^. A convenient alternative to SNP-based genotyping is offered by the commercial software SeqSphere, which implements a core genome multilocus sequence typing scheme (cgMLST) for the analysis of genomic diversity in *C. difficile* ^18^ and other organisms. Indeed, cgMLST ^18^ confirmed the prior conclusion from genomic SNP analyses ^19^ that a common clone of *C. difficile* had been isolated over two successive years at a hospital in China. However, we are not aware of a quantitative comparison of the sensitivity of both methods.

cgMLST of genomic sequences of a variety of bacterial pathogens can also be performed with EnteroBase (http://enterobase.warwick.ac.uk/), which has been developed over the last few years with the goal of facilitating genomic analyses by microbiologists ^20^. EnteroBase automatically retrieves Illumina short-read sequences from public short-read archives. It uses a consistent assembly pipeline to automatically assemble these short-reads into draft genomes consisting of multiple contigs, and presents the assembled genomes together with their metadata for public access ^21^. It also performs the same service on sequencing data uploaded by its registered users. Assembled genomes that pass quality control are genotyped by MLST at the levels of 7-gene MLST, ribosomal MLST (rMLST), core-genome MLST (cgMLST) and whole-genome MLST (wgMLST) ^20,21^. EnteroBase supports subsequent analyses based on either SNPs or cgMLST alleles using the GrapeTree or Dendrogram visualisation tools ^22^. EnteroBase also assigns these genotypes to populations by hierarchical clustering (HierCC), which permits finding close relatives at the global level ^21^. Originally, EnteroBase was restricted to the bacterial genera *Salmonella, Escherichia, Yersinia*, and *Moraxella* but since January, 2018, EnteroBase has included a database for genomes and their metadata for the genus *Clostridioides*. In March, 2019, EnteroBase contained 13,515 draft genomes of *C. difficile* plus one genome of *C. mangenotii*. These included over 900 unpublished draft genomes that were sequenced at the Leibniz Institute DSMZ, as well as 80 complete genome sequences based on Pacific Biosciences plus Illumina sequencing technologies. It also included 862 unpublished draft genomes that were sequenced at the Wellcome Sanger Institute.

Here we show that EnteroBase cgMLST and SNP analyses attain comparable levels of resolution. We also summarise the genomic diversity that accumulated during recurring infections within single patients as well as transmission chains within individual hospitals and between neighbouring hospitals in Germany, and show that it can be detected by HierCC. We also demonstrate that HierCC can be used to identify bacterial populations at various epidemiological levels ranging from recent transmission chains through to pandemic and endemic spread, and relate these HierCC clusters to genotypes that were previously identified by PCR ribotyping and 7-gene MLST. These observations indicate that cgMLST and HierCC within EnteroBase can provide a common language for communications and interactions by the global community who is combatting disease caused by *C. difficile.*

## RESULTS

### Implementation of MLST schemes in EnteroBase

cgMLST in EnteroBase consists of a defined subset of genes within a whole-genome MLST scheme that represents all single copy orthologs within the pan-genome of a representative set of bacterial isolates. To this end, we assembled the draft genomes of 5,232 isolates of *C. difficile* from public short-read archives, and assigned them to ribosomal sequence types (rSTs) according to rMLST, which indexes diversity at 53 loci encoding ribosomal protein subunits on the basis of existing exemplar alleles at PubMLST ^23^. We then created a reference set of 442 genomes consisting of one genome of *C. mangenotii* ^1^, 18 complete genomes from GenBank, 81 closed genomes from our work, and the draft genome with the smallest number of contigs from each of the 343 rSTs (https://tinyurl.com/Cdiff-ref). The *Clostridioides* pan-genome was calculated as described ^21^, and used to define a wgMLST scheme consisting of 13,763 genetic loci (http://enterobase.warwick.ac.uk/species/clostridium/download_data). EnteroBase uses the wgMLST scheme to call loci and alleles from each assembly, and extracts the allelic assignments for the subsets corresponding to cgMLST, rMLST and 7-gene MLST from those allelic calls. The cgMLST subset consists of 2,556 core genes which were present in ≥98% of the reference set, intact in ≥94% and were not excessively divergent (Figure 1).

**Figure 1.**
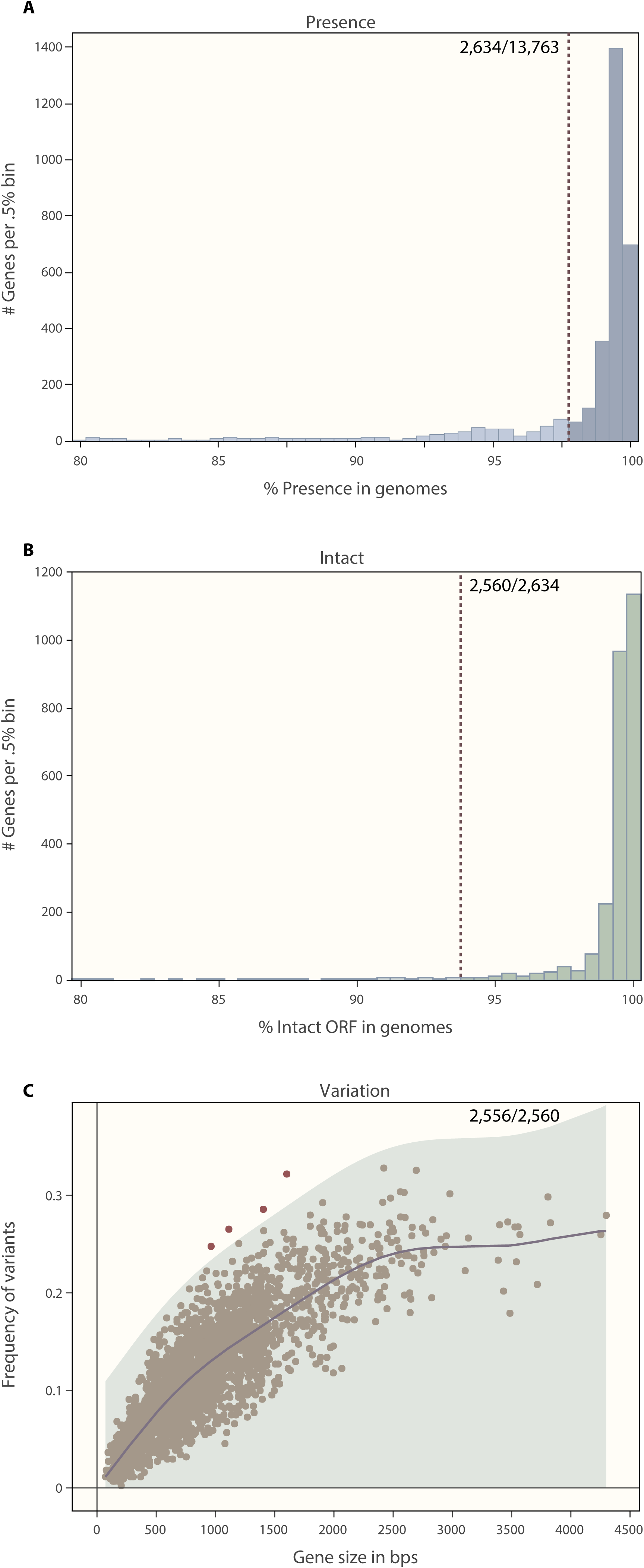
Criteria for inclusion in a cgMLST scheme of a subset of wgMLST genes based on their properties in a reference set of 442 genomes (https://tinyurl.com/Cdiff-ref). (A) Numbers of genes *versus* frequency (% presence) within the reference set. 2,634 genes satisfied the cut-off criterion of ≥98% presence (dashed line). (B) Numbers of genes *versus* intact open reading frame (% intact ORF) within the 2,634 genes from A. 2,560 genes satisfied the cut-off criterion of ≥94% intact ORF (dashed line). (C) Frequency of allelic variants versus gene size among the 2,560 genes from B. The genetic diversity was calculated using the GaussianProcessRegressor function in the sklearn module in Python. This function calculates the Gaussian process regression of the frequency of genetic variants on gene sizes, using a linear combination of a radial basis function kernel (RBF) and a white kernel ^48^. The shadowed region shows a single-tailed 99.9% confidence interval (≤3 sigma) of the prediction. 2,556 loci fell within this area and were retained for the cgMLST scheme, while four were excluded due to excessive numbers of alleles.

### Suitability of cgMLST for epidemiological analyses of transmission chains

Multiple bacterial strains from local outbreaks of *C. difficile* disease in hospitals in Oxfordshire and China differ by very few SNPs ^13,15,19^. The published data indicates a 95% probability that the genomes of pairs of strains from a transmission chain differ by two non-recombinant SNPs, or less ^13^. Bletz *et al.* ^18^ have concluded that sets of strains from transmission chains can also be identified because they differ by ≤6 cgMLST allelic differences in pairwise comparisons. To test this conclusion, we performed a quantitative comparison between the numbers of cgMLST allelic differences and the numbers of non-recombinant SNPs in isolates from multiple epidemiological chains (Figure 2). We calculated the intra-patient pairwise genetic distances by both measures between 176 isolates from four patients with recurring CDI (*C. difficile* infection). We also analysed the intra-outbreak distances with 63 isolates from four transmission chains in multiple hospitals. We also re-examined the pairwise distances from the comprehensive sample of 1,158 isolates collected over several years in four hospitals in Oxfordshire, UK ^13^. All three analyses yielded a strong linear relationship (R^2^, 0.71-0.93) between the numbers of different cgMLST alleles and the numbers of non-recombinant SNPs (Figure 2). The slope of the regression lines was close to 1.0, indicating a 1:1 increase in cgMLST allelic differences with numbers of SNPs. The same data were also investigated with cgMLST calculated with the commercial program SeqSphere ^18^, and yielded similar results except that the slope was lower due to lower discriminatory power of the SeqSphere cgMLST scheme (lower panels in Figure 2).

**Figure 2.**
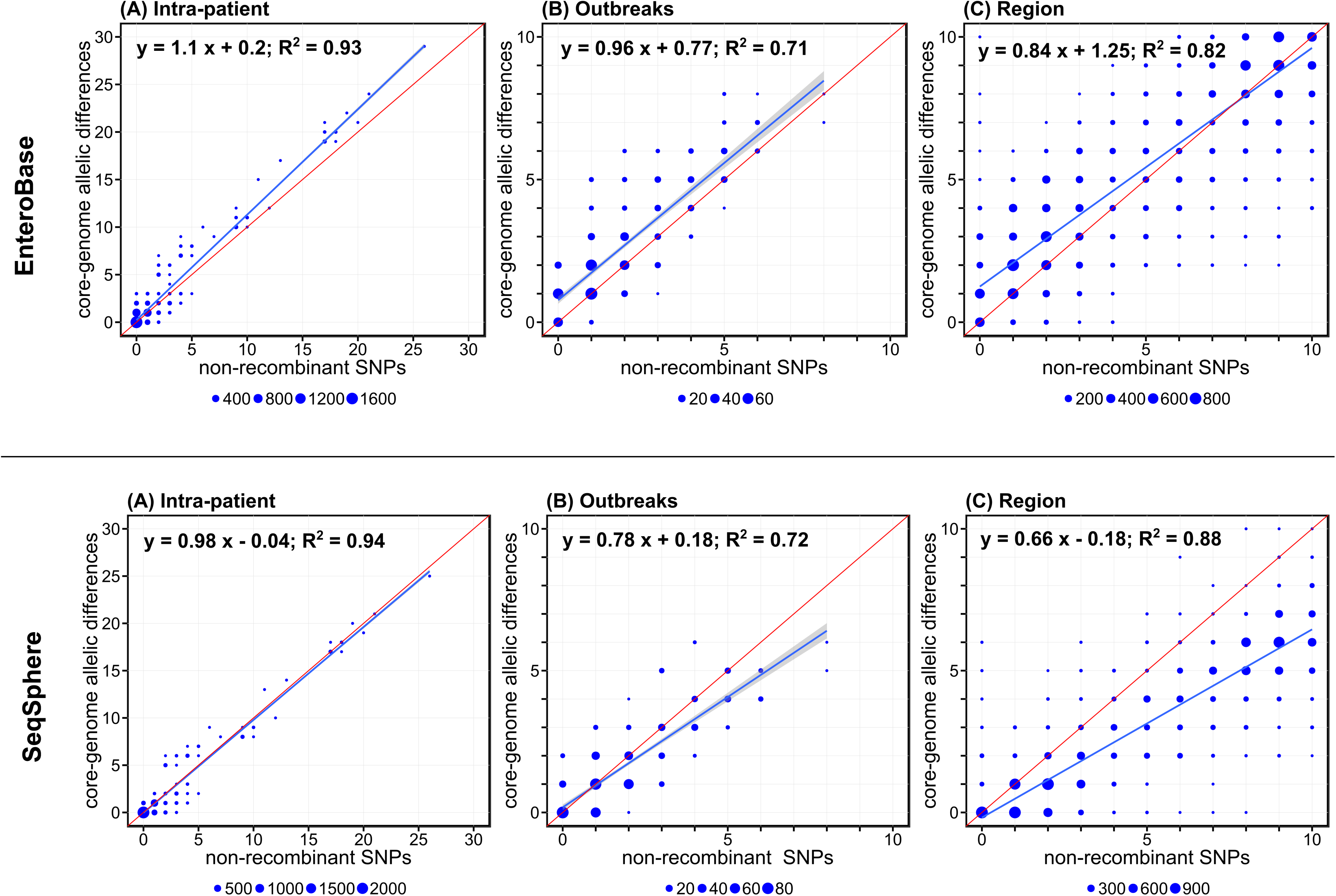
Plots of genomic distances from cgMLST versus non-recombinant SNPs. Upper panels are based on EnteroBase cgMLST, lower panels are based on SeqSphere cgMLST ^18^. **(A) Intra-patient diversity.** Pairwise genomic distances among 176 isolates from four patients, each of which suffered from two episodes of recurrent CDI, 80 to 153 days apart. **(B) Intra-outbreak diversity.** Pairwise genomic distances among isolates from four previously reported CDI outbreaks, including an outbreak protracting over two years in a hospital in China (number of isolates, n, 12) 19, an outbreak involving two hospitals in Southern Germany (n = 9) ^28^, and two transmission chains with ribotypes 027 (n=22) and 106/500 (n=20) in a hospital in Madrid, Spain ^14^. **(C) Regional diversity.** Pairwise genomic distances among isolates from a comprehensive sample of 1,158 isolates collected from CDI patients in several hospitals in Oxfordshire, UK, between 2007 and 2011 ^13^. Distances up to 10 SNPs / 10 cgMLST allelic differences are plotted; the correlation of larger genomic distances is shown in Suppl. Figure 8.

Eyre *et al.* ^13^ concluded that it is possible to recognize genomes that reflect direct transmission between two hospital patients because they tend to differ by two SNPs or less. Our analylsis of the Oxfordshire dataset indicated that they would also have been recognized by cgMLST in EnteroBase because genomes which differed by two cgMLST alleles usually also differed by ≤2 SNPs according to a binary logistic regression model (*p* = 89%; 95% confidence interval, 88-89%) (Figure 3). We also compared the genetic distances between 242 genomes from Oxfordshire which had been isolated during the initial six month trial and 916 genomes during the subsequent three years (April 2008 to March 2011)^13^. Based on cgMLST, 35% of the latter genomes (318/916) also differed at ≤2 core-genome alleles from any earlier genome in that study, and 34% (316/916) differed by ≤2 SNPs. The genomes identified by both methods were also largely concordant (89% matching entries). Thus, our results confirm that cgMLST can be used for epidemiological investigations with similar results to SNP analysis.

**Figure 3.**
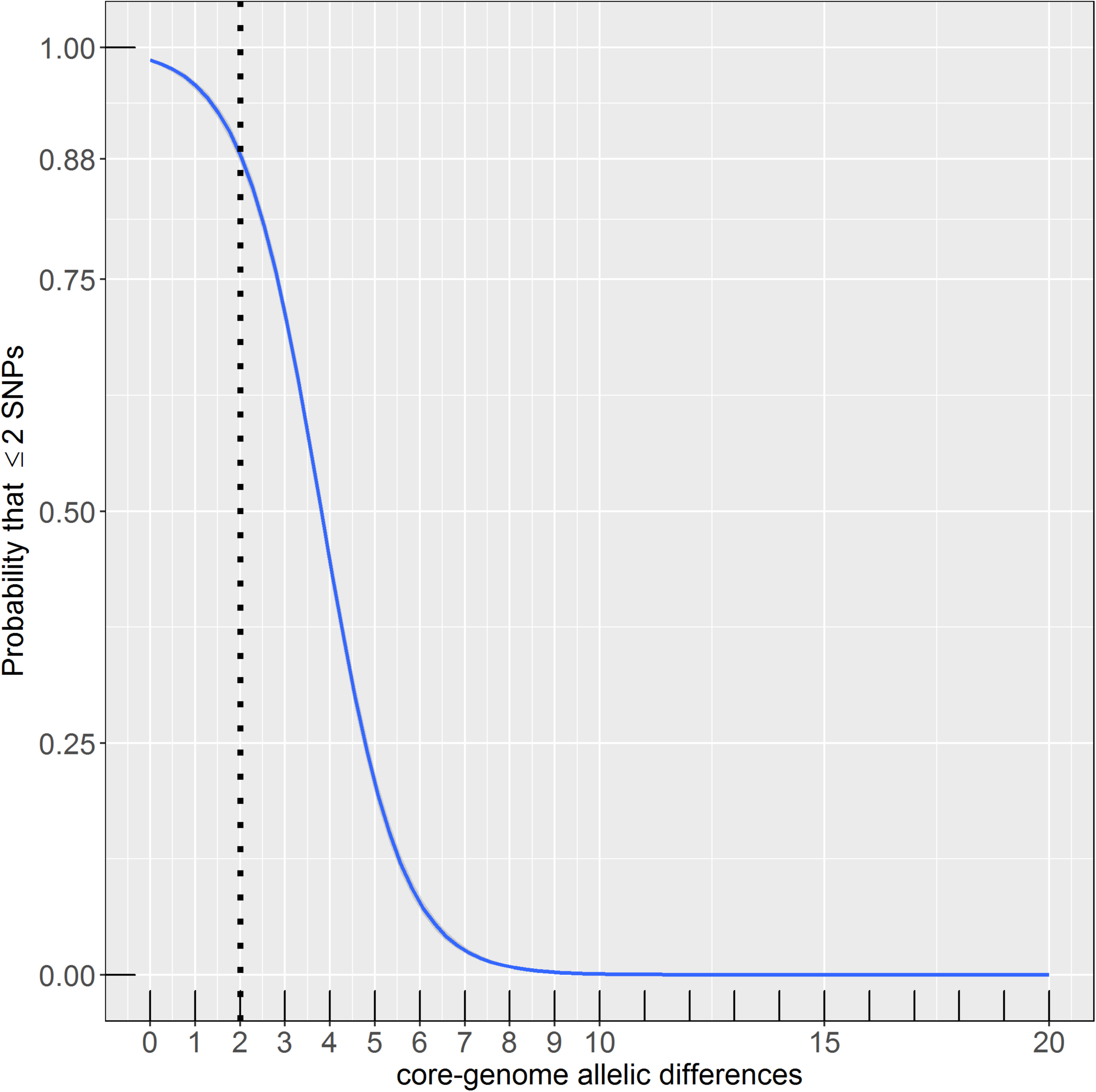
Binary logistic regression model to determine the probability that two genomes are related at ≤2 SNPs, given a certain difference in their cgMLST allelic profiles, based on the Oxfordshire dataset ^13^. The number of SNPs was encoded as a binary dependent variable (1 if ≤2 SNPs, 0 if otherwise) and the number of allelic differences was used as a predictor variable.

### Local and regional spread

SNP analyses are computer intensive and are only feasible with limited numbers of genomes ^24^. cgMLST can be used to analyse up to 100,000 genomes with GrapeTree, but once again analyses involving more than 10,000 genomes are very computer intensive ^22^. EnteroBase has therefore implemented single-linkage hierarchical clustering (HierCC) of cgMLST data at multiple levels of relationship after excluding missing data in pairwise comparisons ^21^ (hierarchical clusters HC0: clusters of indistinguishable core-genome Sequence Types (cgSTs); HC2: clusters with pairwise distances of up to two cgMLST alleles; HC5; HC10; HC20; HC50; … HC2500). Here we address the nature of the genetic relationships that are associated with multiple levels of HierCC clustering within 13,515 publicly available *C. difficile* genomes, and examine which levels of pairwise allelic distances correspond to epidemic outbreaks and to endemic populations, respectively.

As described above, multiple isolates from individual patients fell into patient-specific HC2 clusters, even when sampled from multiple episodes of recurrent CDI that had occurred 80 to 153 days apart (Suppl. Table S1) (patients D, F, and G in Figure 4). In these cases, relapsing disease likely reflected continued colonization after initially successful therapy. However, multiple HC2 clusters were also identified. Some isolates from patient F differed by 12-21 cgMLST allelic differences from the bulk population (Figure 4). Such strain diversity likely reflects co-infections with multiple related strains. In patient E, the two CDI episodes differed by >2,000 allelic differences (Figure 4), indicating an independent reinfection with an unrelated strain. Hence, discrimination between relapse and reinfection based on cgMLST appears to be straightforward, although relapse might be simulated on occasion by reinfection with identical strains because the environment around CDI patients tends to be contaminated with *C. difficile* spores ^25^. We note that time intervals between the relapse episodes investigated here had been between 16 and 22 weeks, beyond the currently recommended threshold of eight weeks for surveillance-based detection of CDI relapses ^26^. Additional similar observations would lend support to extending this definition to a longer timeframe ^27^.

**Figure 4.**
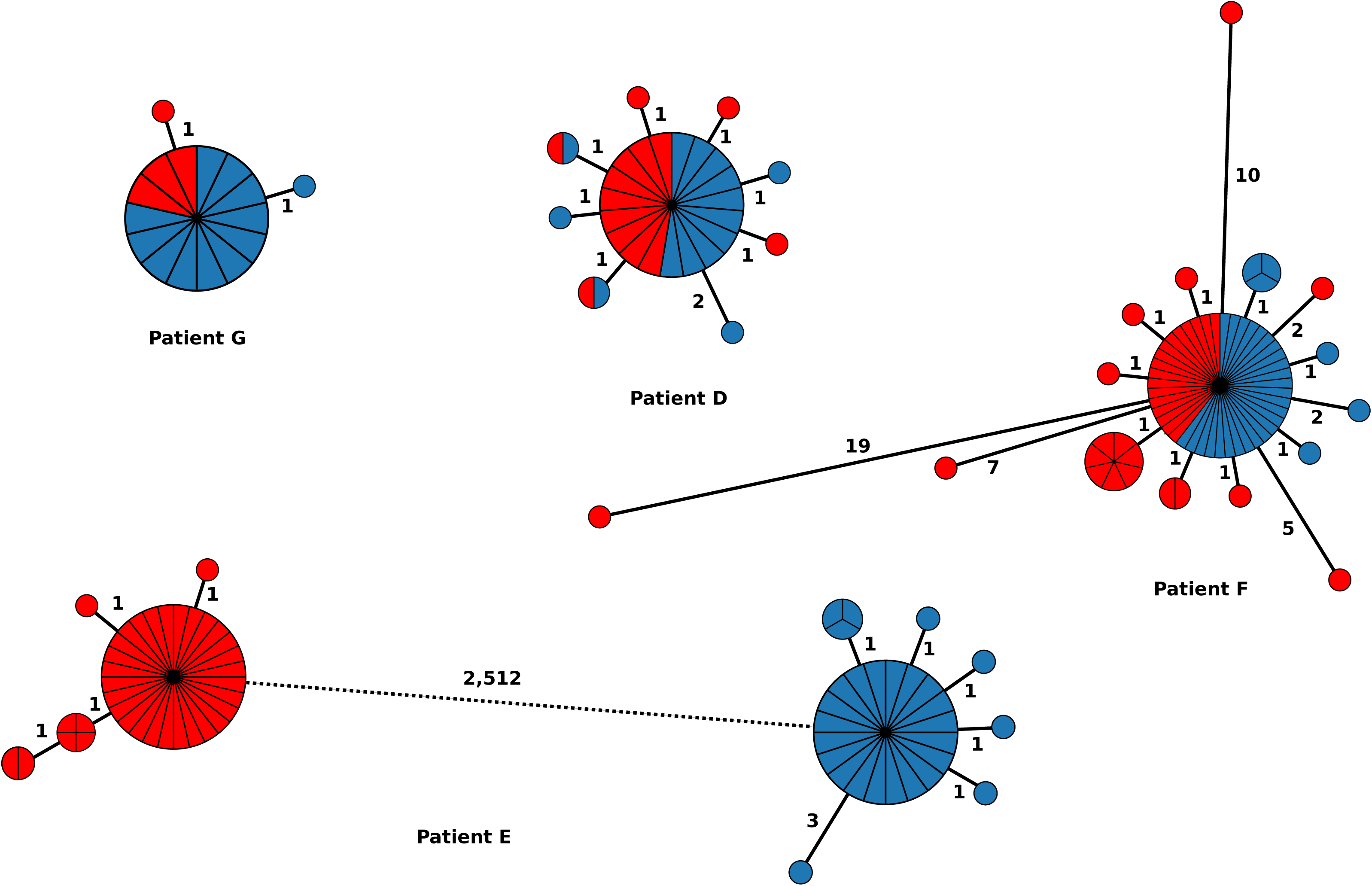
Minimum-spanning trees indicating the population structure of *C. difficile* in four patients with recurrent CDI episodes. Red, first episode; blue, second episode.

Although isolates from a single HC2 cluster are likely to represent an outbreak, it is readily conceivable that multiple HC2 clusters might be isolated from a single epidemiological outbreak due to accumulation of genetic diversity over time or incomplete sampling which resulted in the absence of missing links. Indeed, the majority of isolates from densely sampled, local outbreaks were related at the HC2 level, but some outbreaks investigated here consisted of more than one HC2 cluster (Figure 5). For example, nine isolates from a recently reported outbreak of ribotype 018 in Germany ^28^ formed four related HC2 clusters, and outbreaks with ribotypes 027 and 106 in a hospital in Spain ^14^ were each affiliated with two or three HC2 clusters (Figure 5). Incomplete sampling of outbreaks may be common because asymptomatic patients may constitute an important reservoir for transmission but are only rarely examined for colonization with *C. difficile* ^29-31^.

**Figure 5.**
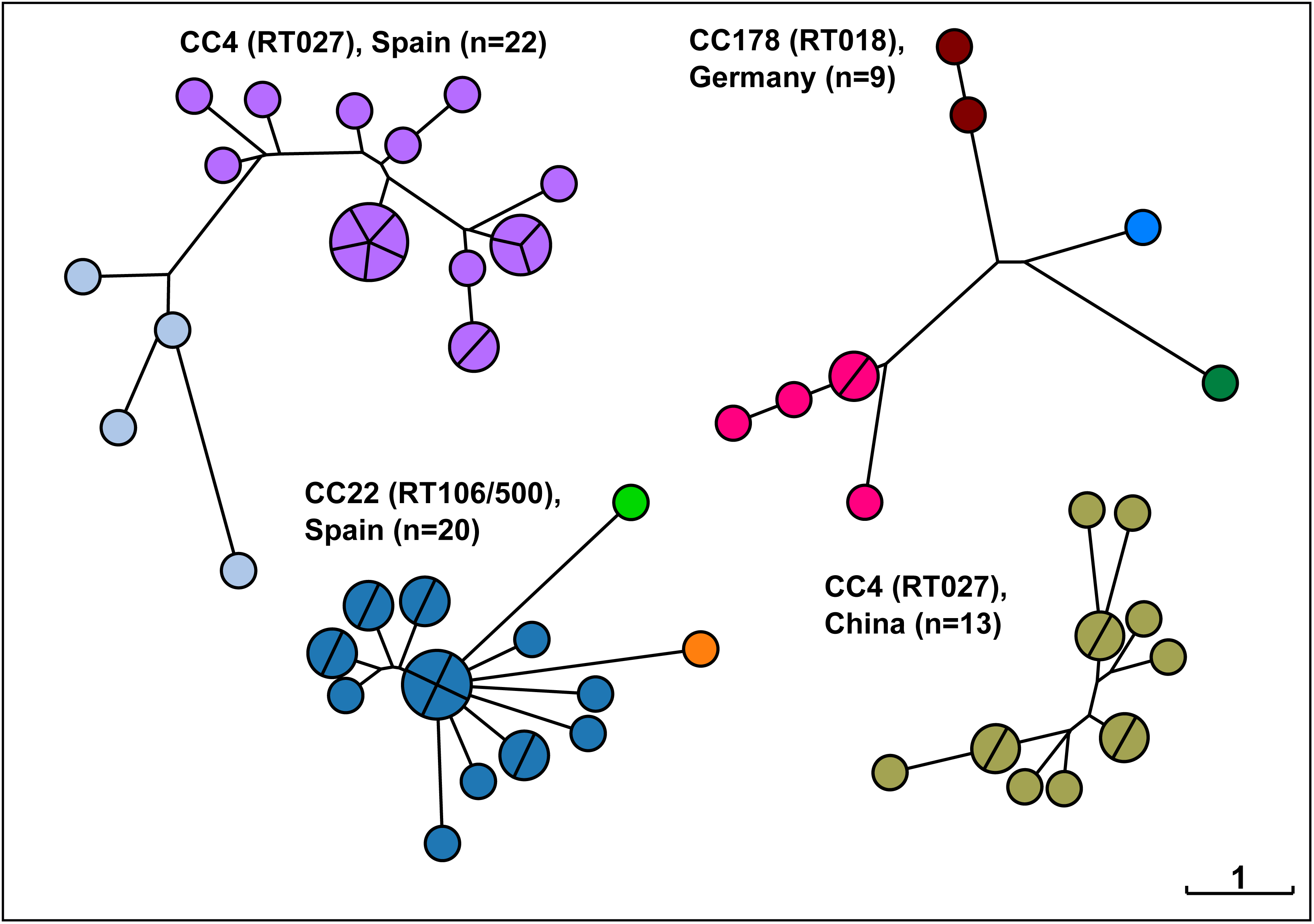
Neighbour-joining trees based on cgMLST showing the phylogenetic relationships among *C. difficile* isolates from previously published CDI outbreaks as indicated ^14,19,28^. Nodes are coloured by HC2. CC, cgST complex; RT, PCR ribotype. The scale, indicating one allelic difference, applies to all trees.

We identified 23 HC2 clusters (encompassing 133 genome sequences) in a dataset of 309 *C. difficile* genome sequences collected from CDI patients in six neighboring hospitals in Germany. These HC2 clusters were associated with individual hospitals (Χ2, p=8.6×10^-5^; Shannon entropy, p=4.2×10^-5^) and even with single wards in these hospitals (Χ2, p=0.01; Shannon entropy, p=6.2×10^-3^). We investigated whether these HC2 clusters reflected the local spread of *C. difficile* within institutions by retrospective analyses of patient location data. Sixty six cases (50%) revealed ward contacts with another patient with the same HC2 cluster (median time interval between ward occupancy: 63 days; range, 0 to 521 days). These results are consistent with the direct transmission for *C. difficile* isolates of the same HC2 cluster (Figure 6). In those cases where the shared ward contacts were separated in time, transmission may have occurred indirectly or from a common reservoir, possibly through environmental spore contamination or through asymptomatically colonized patients^14,29,30^. We also detected 15 HC2 clusters that included isolates from two or more hospitals in the region. Subsequent analyses of patient location data confirmed that some of these HC2 clusters were directly associated with patient transferrals between the hospitals (Figure 6). Hence, hierarchical clustering of *C. difficile* genome sequences in conjunction with retrospective analysis of patient movements revealed multiple likely nosocomial transmission events, none of which had been detected previously based on routine surveillance.

**Figure 6.**
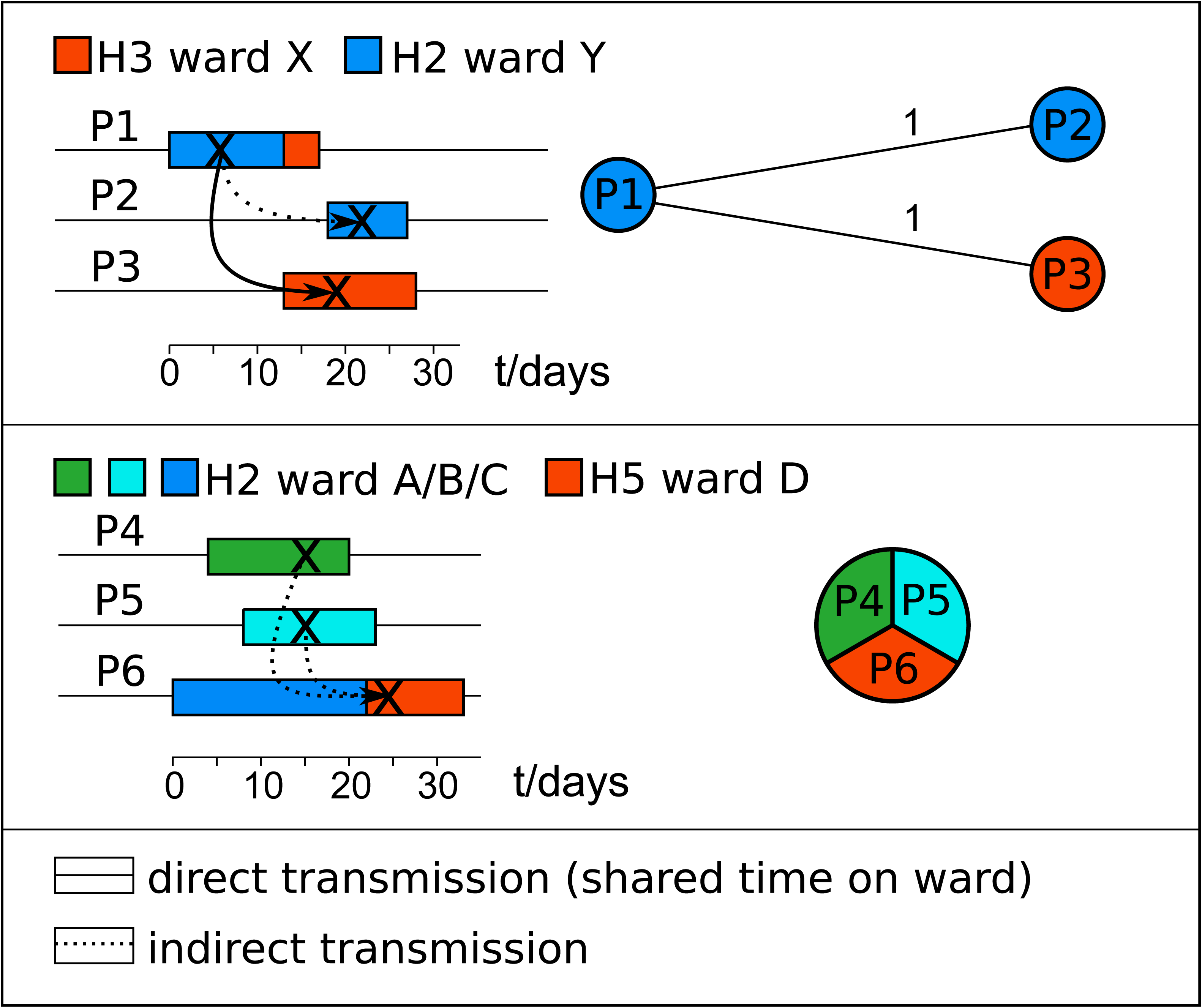
Timelines of two transmission chains, discovered retrospectively through inspection of files from CDI patients with closely related *C. difficile* isolates (HC2). Colours indicate hospital wards, ‘X’ indicate diagnosis of CDI, and arrows indicate presumed transmission pathways. Minimum-spanning trees indicating genomic distances among *C. difficile* isolates are shown on the right. Upper panel: patient P1 was diagnosed with CDI in hospital H3 and transferred to hospital H3 15 days later. Another five and six days later, respectively, patients P2 in hospital H2 and P3 in hospital H3 got diagnosed with CDI with closely related strains. Both these patients were on the same wards as the initial patient, who probably had been the source for the pathogen. Since there was no temporal overlap between these patients in hospital H2, transmission may have occurred indirectly, possibly through environmental contamination. Lower panel: another putative transmission chain involved three patients that had shared time in hospital H2. Patients P4 and P5 got diagnosed with CDI on the same day after they had shared seven days in this hospital, albeit on different medical wards. The third patient developed CDI with the same *C. difficile* cgST in a different hospital (H5), but had previously stayed at hospital H2 during the time when CDI got diagnosed in the first two patients.

### Pandemic strains and endemic populations

Pandemic spread of *C. difficile* over up to 25 years has been inferred previously on the basis of molecular epidemiology with other, lower resolution techniques ^32^. The majority of isolates in EnteroBase that are known to representing such pandemic strains were related at the level of HC10, including epidemic ribotype 017 (cluster HC10|17) ^11^, each of two previously reported fluoroquinolone-resistant lineages of ribotype 027 (HC10|4, HC10|9) ^9^, or livestock-associated ribotype 078/126 (HC10|1) ^33^ (Suppl. Figure 1).

Endemic populations have also been described by ribotyping and phylogenetic analyses, some of which have formed a source for the emergence of epidemic or pandemic strains ^2,9^. Many such endemic populations may be represented by HC150 clusters. Clustering at HC150 was well supported statistically (Suppl. Figure 2), and the frequency distribution of pairwise genomic distances indicated that multiple database entries clustered at <150 cgMLST allelic differences (Suppl. Figure 3). A cgMLST-based phylogenetic tree of 13,515 *C. difficile* genomes showed 201 well separated HC150 clusters, each encompassing a set of closely related isolates, plus 209 singletons (Figure 7). Because these HC150 clusters are based on cgMLST genetic distances, we will refer to them as ‘cgST complexes’, abbreviated as CCs. Genomes from each of the major CCs have been collected over many years in multiple countries, indicating their long-term persistence over wide geographic ranges (Table 1).

**Table 1.** Characteristics of cgST complexes (CC) with ≥100 entries.

| CC<br>(HC150) | PCR Ribotype | Number of<br>entries | Sampling<br>years | Number of<br>countries | % isolates<br>in HC2>2 <sup>1</sup> | % isolates<br>from animal<br>hosts |
| --- | --- | --- | --- | --- | --- | --- |
| 4 | 027 | 2,669 | 1985-2018 | 27 | 77 | 0 |
| 1 | 078, 126 | 1,222 | 1994-2018 | 26 | 61 | 17 |
| 17 | 017 | 769 | 1990-2017 | 24 | 64 | 0 |
| 3 | 001 | 768 | 1980-2017 | 16 | 62 | 0 |
| 6 | 020, 404 | 768 | 1995-2017 | 14 | 43 | 1 |
| 2 | 002 | 702 | 2006-2017 | 15 | 51 | 1 |
| 22 | 106, 500 | 531 | 1997-2017 | 7 | 59 | 3 |
| 86 | 005 | 468 | 1980-2017 | 8 | 41 | 0 |
| 34 | 014 | 421 | 1995-2017 | 10 | 35 | 0 |
| 55 | 015 | 318 | 2006-2017 | 6 | 37 | 0 |
| 71 | 014, 020 | 315 | 2004-2017 | 16 | 40 | 1 |
| 145 | 015 | 284 | 2006-2016 | 7 | 39 | 0 |
| 256 | 023 | 268 | 2001-2015 | 6 | 40 | 0 |
| 79 | 010 | 249 | 2003-2018 | 7 | 53 | 3 |
| 178 | 018, 356 | 243 | 2006-2017 | 7 | 52 | 0 |
| 242 | 039 | 199 | 2008-2017 | 4 | 58 | 1 |
| 10 | 012 | 159 | 1996-2017 | 7 | 52 | 0 |
| 88 | 014 | 132 | 1996-2016 | 9 | 33 | 8 |
| 11 | 070 | 110 | 2006-2017 | 6 | 32 | 0 |
| 187 | 054 | 109 | 2007-2018 | 6 | 47 | 0 |
| 141 | 001, 026 | 107 | 2007-2016 | 2 | 7 | 0 |
| 391 | 081 | 105 | 1996-2016 | 4 | 31 | 0 |
| 49 | 011, 056, 446 | 103 | 2001-2017 | 5 | 35 | 0 |

<sup>1</sup> isolates in HC2 clusters with >2 entries

**Figure 7.**
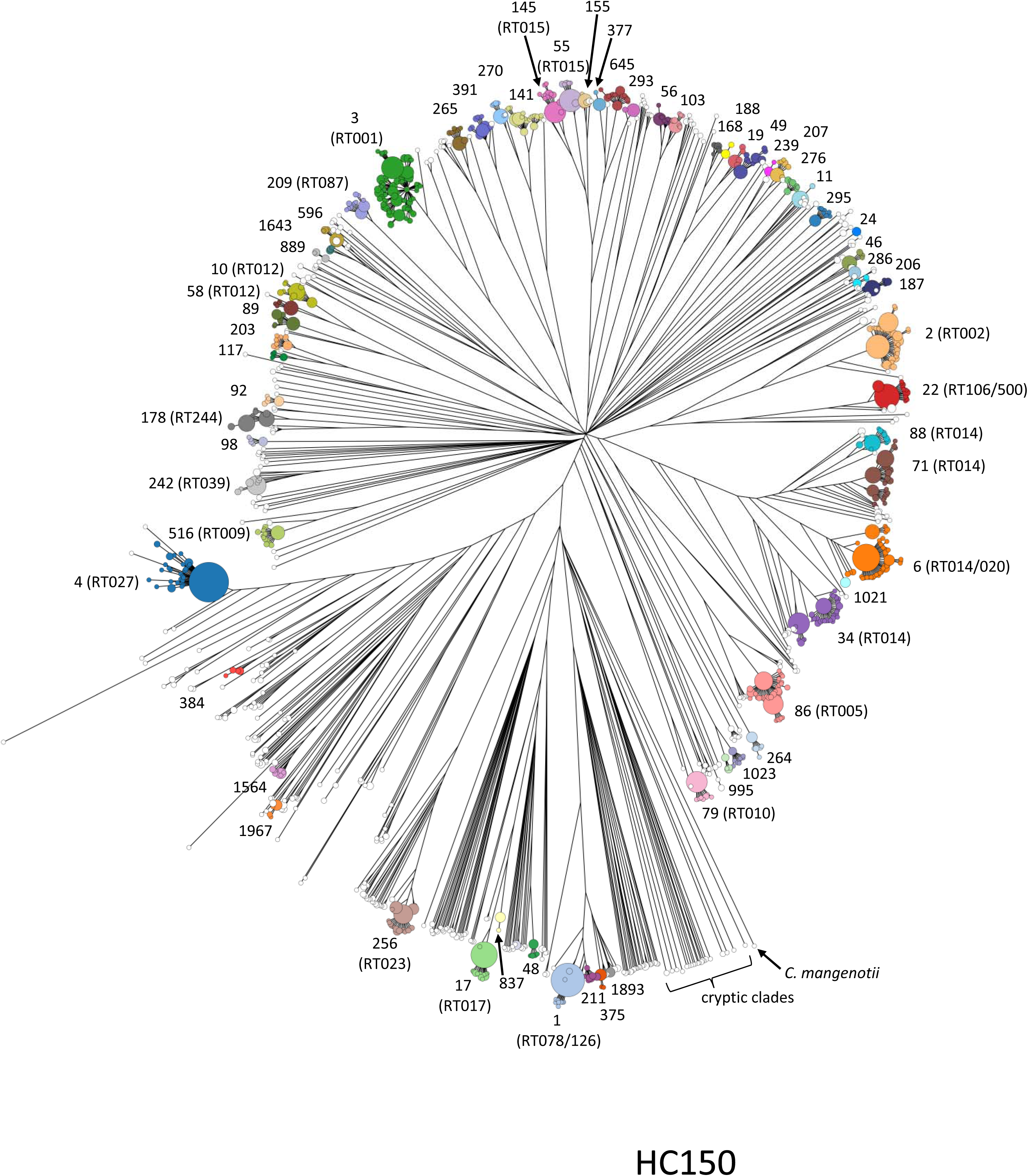
Rapid-neighbour-joining phylogenetic tree based on cgMLST variation from 13,515 *C. difficile* genomes. Colours and numerals indicate CCs (HC150 clusters) with ≥10 entries, and information on predominant PCR ribotypes is provided in brackets.

We compared HC150 clustering with PCR ribotyping for 2,263 genomes spanning 84 PCR ribotypes for which PCR ribotyping data were available in EnteroBase. These included 905 genomes which we ribotyped ourselves (Suppl. Table S2) as well as several hundred genomes for which we had manually retrieved ribotype information from published data. The correlation between HC150 clustering and ribotyping was high (adjusted Rand coefficient, 0.92; 95% confidence interval, 0.90-0.93). However, our analysis also revealed that PCR ribotypes did not always correspond to phylogenetically coherent groupings. PCR ribotypes 002, 015 and 018 were each distributed across multiple phylogenetic branches (Figure 8). Furthermore, multiple ribotypes were assigned to genomes with indistinguishable cgMLST alleles. These included ribotypes 106 and 500, ribotype 014 and 020, and ribotypes 066, 126, 413 and 078, as well as other examples in Figure 8. In contrast, HC150 clusters corresponded to clear-cut phylogenetic groupings within a phylogenetic tree of core genes (Figure 8-HC150).

**Figure 8.**
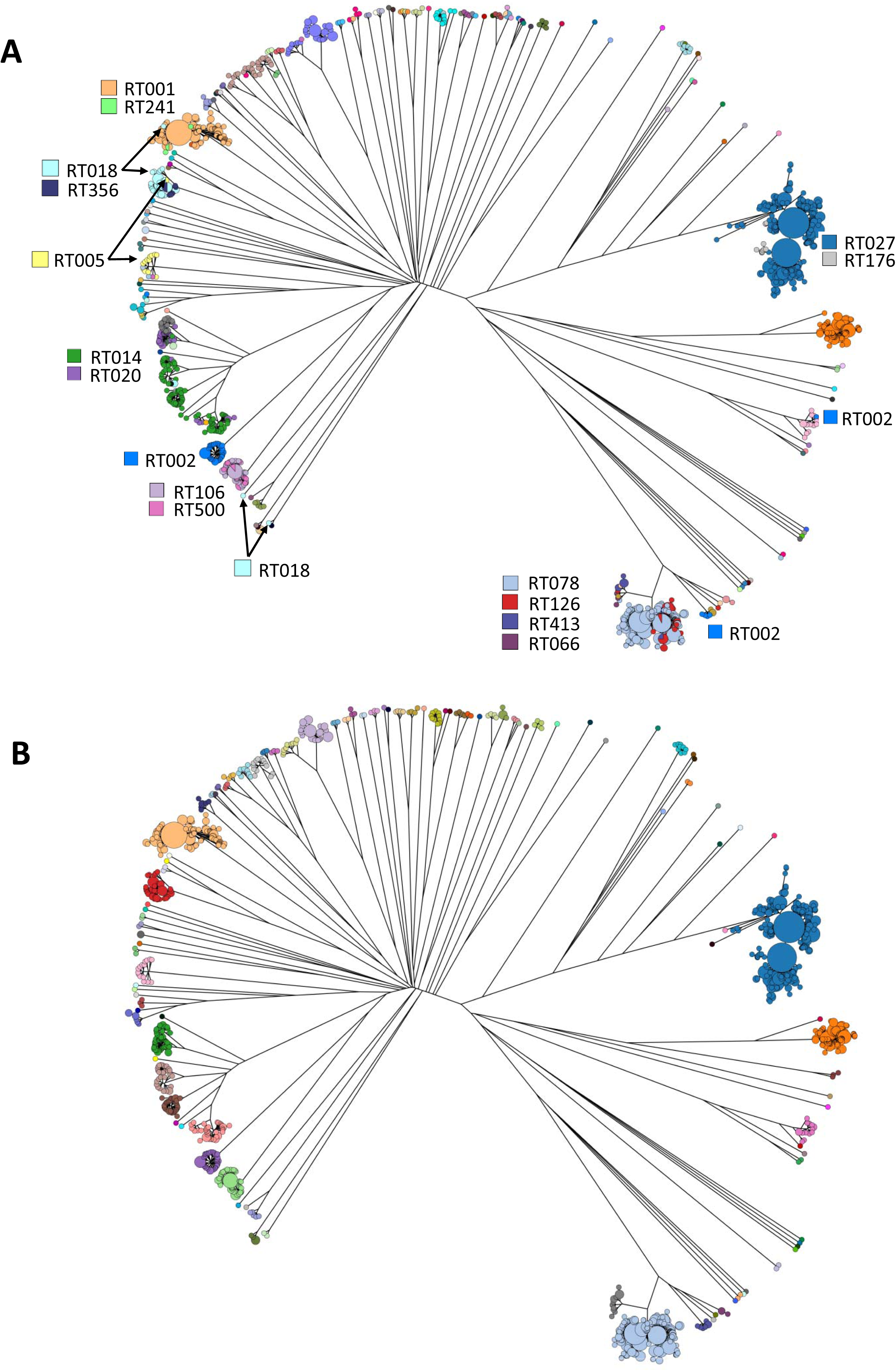
Rapid-neighbour-joining phylogenetic tree based on cgMLST variation from 2,263 *C. difficile* genomes, for which PCR ribotyping information is available. (A) Nodes are coloured by PCR ribotype as indicated. (B) Nodes are coloured by CC.

Rarefaction analysis indicated that the currently available genome sequences represent about two-thirds of extant CC diversity (Suppl. Figure 4). It is not clear why so many populations of *C. difficile* co-exist, but at least some of this diversity may be due to the occupation of distinct ecological niches, as exemplified by differential propensities for colonizing non-human host species (Table 1) ^34,35^. Furthermore, the different CCs may differ in their aptitudes for epidemic spread, as suggested by very different proportion of genomes assigned to HC2 chains: compare CC141 with 7% in HC2 clusters *versus* CC4 with 77% (Table 1).

### Higher population levels

HierCC can identify clusters at higher taxonomic levels ^21^. In *C. difficile*, HC950 clusters seem to correspond to deep evolutionary branches (Suppl. Figure 5) and HC2000 clusters were congruent with the major clades reported previously ^36^, except that cluster HC2000|2 encompassed clade 1 plus clade 2 (Suppl. Figure 6). Finally, HC2500 may correspond to the subspecies level, because it distinguished between *C. difficile* and distantly related “cryptic clades” (Suppl. Figure 7).

## DISCUSSION

In this study, we demonstrate that the application of cgMLST to investigations of local *C. difficile* epidemiology yields results that are quantitatively equivalent to those from SNP analyses. This is a major advance because SNP analyses require specific bioinformatic skills and infrastructure, are time consuming, and not easily standardized ^16^. Even though a cgMLST scheme for *C. difficile* had been published recently, its ability to identify closely related isolates and the inferred genomic distances had not been compared to SNP analyses in any quantitative way ^18^. In EnteroBase, cgMLST is based on de-novo assembly of sequencing reads, which is also known to introduce errors associated with the assembler algorithms. However, EnteroBase uses Pilon ^37^ to polish the assembled scaffolds and evaluate the reliability of consensus bases of the scaffolds, thereby achieving comparable accuracy to mapping-based SNP analyses. When applied to a large dataset of *C. difficile* genomes from hospital patients in the Oxfordshire region (UK), cgMLST and SNP analysis were largely consistent (89% match) at discriminating between isolates that were sufficiently closely related to have arisen during transmissions chains from others that were epidemiologically unrelated. Similarly, cgMLST identified numerous HC2 clusters of strains in *C. difficile* isolates that might have arisen during transmission chains in six neighboring hospitals in Germany. These assignments were in part consistent with retrospective investigation of patient location data, although none of the outbreaks had been detected previously by standard epidemiological surveillance by skilled clinical microbiologists. Recent publications propose that prospective genome sequencing of nosocomial pathogens should be applied routinely at the hospital level to guide epidemiological surveillance ^38^. Our data indicates that the combination of genome sequencing with cgMLST may identify nosocomial transmission routes of *C. difficile* more effectively than presently common practice, and hence could help to reduce pathogen spread and the burden of disease.

Infectious disease epidemiologists frequently seek to know if new isolates of bacterial pathogens are closely related to others from elsewhere, i. e. if they are part of a widespread outbreak. Unlike a previous cgMLST implementation ^18^, EnteroBase supports this goal by taking full advantage of rapidly growing, public repositories of short-read genome sequences ^21^. In contrast to short-read archives, however, where stored sequence data are not readily interpretable without specialized bioinformatic tools ^39^, EnteroBase enables contextual interpretation of a growing collection (13,515 entries as of March 2019) of assembled, quality-controlled *C. difficile* genome sequences and their associated metadata. Importantly, phylogenetic trees based on cgMLST allelic profiles from many thousand bacterial genomes can be reconstructed within a few minutes, whereas such calculations currently are prohibitively slow based on SNP alignments ^21^. Genome sequencing reads from newly sampled *C. difficile* isolates can be uploaded to EnteroBase and compared to all publicly available genome data within hours, without requiring any command-line skills.

In March 2019, the *Clostridioides* database contained >10,000 cgSTs, indicating that the majority of genomes was unique. Furthermore, after assembly, draft genomes contain missing data and many cgSTs have unique cgST numbers but are identical to other cgSTs, except for missing data. Hence, individual cgST numbers are only rarely informative. However, indistinguishable cgSTs are clustered in common HierCC HC0 clusters, which ignore missing data. Similarly, EnteroBase provides cluster designations at multiple levels of HierCC, enabling rapid identification of all cgSTs that are related at multiple levels of genetic distance. The data presented here shows that HierCC designations can facilitate communications between researchers, clinicians and public-health officials about transmission chains, epidemic outbreaks, endemic populations and higher phylogenetic lineages up to the level of subspecies. HierCC will also enable comparisons to previously published data because we have provided a correspondence table between HC150 clusters and PCR ribotypes. A full understanding of the population structure of *C. difficile* and its relationship to epidemiological patterns will require additional study because many of the clusters described here have not yet been studied or described. However, this task can be addressed by the global community due to the free public access to such an unprecedented amount of genomic data from this important pathogen.

## Methods and Materials

### Sampling

309 *C. difficile* isolates were collected at a diagnostic laboratory providing clinical microbiology services to several hospitals in central Germany. To assemble a representative sample, we included the first 20 isolates from each of six hospitals from each of three consecutive calendar years (Suppl. Table S2). For investigation of recurrent CDI, a set of 183 *C. difficile* isolates were collected in a diagnostic laboratory in Saarland, Germany. Here, primary stool culture agar plates were stored at 4°C for five months to eventually enable the analysis of multiple plates representing episodes of recurrent *C. difficile* infection from individual patients, who had developed recurrent disease by then and could be chosen with hindsight. It was attempted to pick and cultivate as many bacterial colonies from each selected plate as possible, resulting in 6 to 36 isolates per CDI episode. In addition, we sequenced the genomes from 383 isolates that had been characterized by PCR ribotyping previously, including 184 isolates sampled from piglets ^8^, 71 isolates from various hospitals in Germany ^3^, and 108 isolates from stool samples collected from nursery home residents (unpublished; Suppl. Table S2).

### PCR ribotyping

PCR ribotyping was performed as described previously ^40^, applying an ABI Prism 3100 apparatus for capillary electrophoresis and comparing banding patterns to the Webribo database (https://webribo.ages.at/).

### Whole genome sequencing

For Illumina sequencing, genomic DNA was extracted from bacterial isolates by using the DNeasy Blood & Tissue kit (Qiagen), and libraries were prepared as described previously ^41^ and sequenced on an Illumina NextSeq 500 machine using a Mid-Output kit (Illumina) with 300 cycles. For generating complete genome sequences, we applied SMRT long-read sequencing on an RSII instrument (Pacific Biosciences) in combination with Illumina sequencing as reported previously ^41^.

### SNP detection and phylogenetic analysis

Sequencing reads were mapped to the reference genome sequence from C. difficile strain R20291 (sequence accession number FN545816) by using BWA-MEM and sequence variation was detected by applying VarScan2 as reported previously ^41^. Sequence variation likely generated by recombination was detected through analysis with ClonalFrameML ^42^ and removed prior to determination of pairwise sequence distances ^15^ and to construction of maximum-likelihood phylogenetic trees with RAxML (version 8.2.9) ^43^. All genome sequencing data were submitted to the European Nucleotide Archive (www.ebi.ac.uk/ena) under study numbers PRJEB33768, PRJEB33779, PRJEB33780 and PRJEB33782.

### Statistical analyses

To determine the probability that two genomes are related at ≤2 SNPs, given a certain difference in their cgMLST allelic profiles, we inferred a logistic regression model using R (^44^, p. 593-609). Genomic relatedness was encoded as a binary response variable (1 if ≤2 SNPs, 0 if otherwise) and the number of core-genome allelic differences was used as a predictor variable. We applied this model to a dataset of 1,158 genome sequences from a previous study, representing almost all symptomatic CDI patients in Oxfordshire, UK, from 2007 through 2011 ^13^. While that original study had encompassed a slightly larger number of sequences, we restricted our analysis to the data (95%) that had passed quality control as implemented in EnteroBase ^20^. We used the SNP data from Eyre’s report ^13^.

The hierarchical single-linkage clustering of cgMLST sequence types was carried out as described ^21^ for all levels of allelic distances between 0 and 2,556. We searched for stable levels of differentiation by HierCC according to the Silhouette index ^45^, a measure of uniformity of the divergence within clusters. The Silhouette index was calculated based on d^’, a normalized genetic distance between pairs of STs, which was calculated from their allelic distance d as: d^’=1-(1-d)^(1/l), where l is the average length (937 bp) of the genes in the cgMLST scheme.

We further evaluated the “stability” of hierarchical clustering using two other criteria. The Shannon index is a measure of diversity in a given population. The Shannon index drops from nearly 1 in HC0, because most cgSTs are assigned to a unique HC0 cluster, to 0 in HC2500, which assigns all sequence types to one cluster. The gradient of the Shannon index between the two extremes reflects the frequencies of coalescence of multiple clusters at a lower HC level. Thus, the plateaus in the curve correspond to stable hierarchical levels, where the Shannon index does not change dramatically with HC level. We also evaluated the stability of hierarchical clustering by pairwise comparison of the results from different levels based on the normalized mutual Information score ^46^ (Fig. S3).

To estimate concordance between cgMLST-based hierarchical clustering and PCR ribotyping, we calculated the adjusted Rand coefficient ^47^ by using the online tool available at http://www.comparingpartitions.info/. To test statistical associations of HC2 clusters with specific hospitals and hospital wards, respectively, we compared Χ2 values and normalized Shannon entropy values (R package ‘entropy’ v.1.2.1) from contingency tables containing real isolate distributions (Suppl. Table S3) and randomly permuted distributions (n=1,000), by using the non-parametric, two-sided Mann-Whitney U test (R package ‘stats’ v.3.5.0).

## Acknowledgements

We thank Vera Junker, Simone Severitt, Nicole Heyer and Carola Berg for excellent technical assistance, Johannes Sikorski for help with *R* and David Eyre for supplying SNP data from his 2013 paper in tabular format. This work was partially funded by the German Center for Infection Research (DZIF), by the Federal State of Lower Saxony (Niedersächsisches Vorab VWZN2889/3215/3266), by the EU Horizon 2020 programme (grant agreement number 643476), the Wellcome Trust (098051), the UK Medical Research Council (PF451). EnteroBase development was funded by the BBSRC (BB/L020319/1) and the Wellcome Trust (202792/Z/16/Z), and the salary of ZZ was also provided by The Wellcome Trust.

---

**Suppl. Figure 1.** Phylogenetic structure of three *C. difficile* pandemics, each of which has spread for about 25 years ^9,11^. Within each pandemic, the majority of isolates is related at level HC10, as indicated by the colours. CC, cgST complex; RT, PCR ribotype.

**Suppl. Figure 2.** Statistical evaluation of hierarchical single-linkage clustering at all levels. Dashed lines indicate hierarchical levels of maximum cluster stability. **(A)** Silhouette index at all hierarchical levels, plotted in steps of 50 allelic differences. Local maxima indicate levels with cohesive, well separated clusters. **(B)** Shannon index at all hierarchical levels. Plateaus in this curve indicate levels with stable clustering. **(C)** Normalized mutual information score comparing the stability of clusters at different hierarchical levels. Dark areas indicate ranges of stable clustering.

**Suppl. Figure 3.** Frequency distribution of pairwise genomic distances (cgMLST allelic differences) among 13,515 *C. difficile* genomes in EnteroBase. Indicated levels of hierarchical clustering are discussed in the text.

**Suppl. Figure 4.** Rarefaction analysis to estimate numbers of extant HC clusters.

**Suppl. Figure 5.** Rapid-neighbour-joining phylogenetic tree based on cgMLST variation from 13,515 *C. difficile* genomes. Colours indicate HC950 clusters.

**Suppl. Figure 6.** Rapid-neighbour-joining phylogenetic tree based on cgMLST variation from 13,515 *C. difficile* genomes. Colours indicate HC2000 clusters.

**Suppl. Figure 7.** Rapid-neighbour-joining phylogenetic tree based on cgMLST variation from 13,515 *C. difficile* genomes. Colours indicate HC2500 clusters.

**Suppl. Figure 8.** Pairwise genomic distances (cgMLST allelic differences versus non-recombinant SNPs) between 1,158 isolates collected from CDI patients in hospitals in Oxfordshire, UK ^13^.

## Supplementary Tables

**Suppl. Table S1.** Time intervals between episodes of recurrent infections and numbers of isolates per episode.

**Suppl. Table S2.** List of *C. difficile* isolates sequenced in this study.

**Suppl. Table S3.** Distribution of 23 HC2 clusters among wards in six neighboring hospitals in Germany.

