## Supplementary Table S2 for "Global genomic population structure of *Clostridioides difficile*"

**Suppl. Table S2.** Distribution of 23 HC2 clusters among wards in six hospitals in Lower Saxony, Germany.

| HC2 \ Hospitals | 1 | 2 | 3 | 4 | 5 | 6 |
|---|---|---|---|---|---|---|
| **1** | 0 | 0 | 1 | 0 | 2 | 0 |
| **2** | 0 | 0 | 2 | 0 | 0 | 0 |
| **70** | 0 | 0 | 3 | 0 | 0 | 0 |
| **76** | 15 | 13 | 19 | 2 | 9 | 8 |
| **85** | 0 | 0 | 2 | 0 | 0 | 0 |
| **109** | 0 | 0 | 0 | 0 | 1 | 2 |
| **479** | 0 | 0 | 0 | 0 | 2 | 0 |
| **491** | 2 | 0 | 0 | 0 | 0 | 0 |
| **1127** | 0 | 1 | 0 | 1 | 6 | 0 |
| **1131** | 0 | 1 | 1 | 1 | 2 | 0 |
| **1206** | 1 | 3 | 1 | 0 | 0 | 1 |
| **1208** | 1 | 0 | 1 | 0 | 0 | 1 |
| **1210** | 0 | 1 | 0 | 0 | 0 | 1 |
| **1225** | 2 | 0 | 1 | 0 | 0 | 0 |
| **1232** | 0 | 0 | 0 | 0 | 1 | 1 |
| **1242** | 0 | 0 | 0 | 1 | 0 | 1 |
| **1243** | 1 | 0 | 0 | 0 | 1 | 1 |
| **1251** | 0 | 0 | 2 | 0 | 1 | 2 |
| **1267** | 0 | 2 | 1 | 0 | 0 | 0 |
| **4415** | 0 | 2 | 0 | 0 | 0 | 0 |
| **4431** | 0 | 0 | 0 | 2 | 0 | 0 |
| **4808** | 0 | 0 | 0 | 0 | 2 | 0 |
| **4823** | 0 | 2 | 0 | 0 | 0 | 0 |
