## Supplementary Table S3 for "Global genomic population structure of *Clostridioides difficile*"

**Suppl. Table S3.** Recurrent infections.

| patient | Δt (weeks) | no. of isolates, episode 1/2 |
|---|---|---|
| D | 21.0 | 15 / 14 |
| E | 11.4 | 28 / 36 |
| F | 16.0 | 32 / 35 |
| G | 21.9 | 12 / 4 |
