## Supplementary figures and images for "Global genomic population structure of *Clostridioides difficile*"

### Supplementary Figure 1

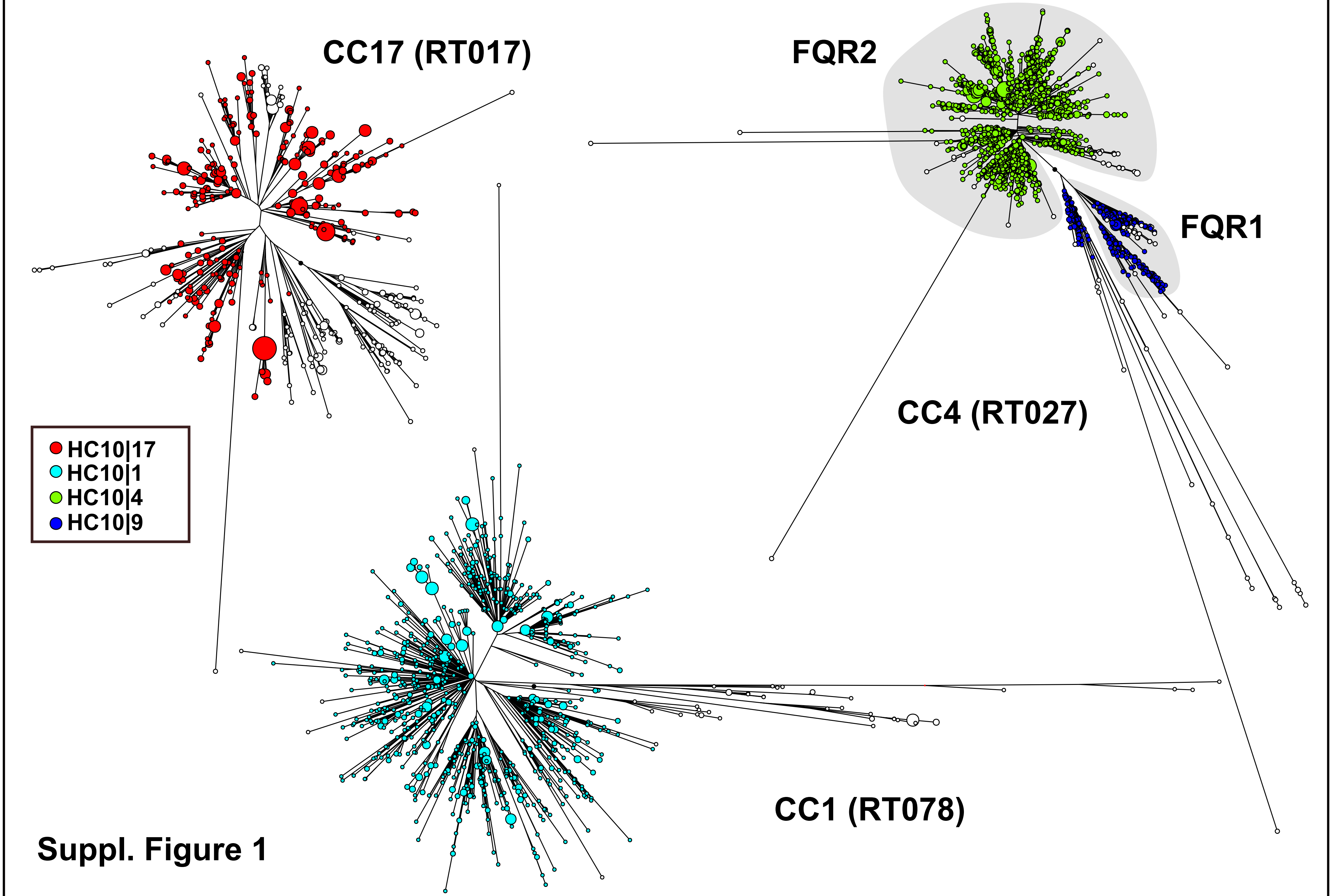


Suppl. Figure 1

### Supplementary Figure 2

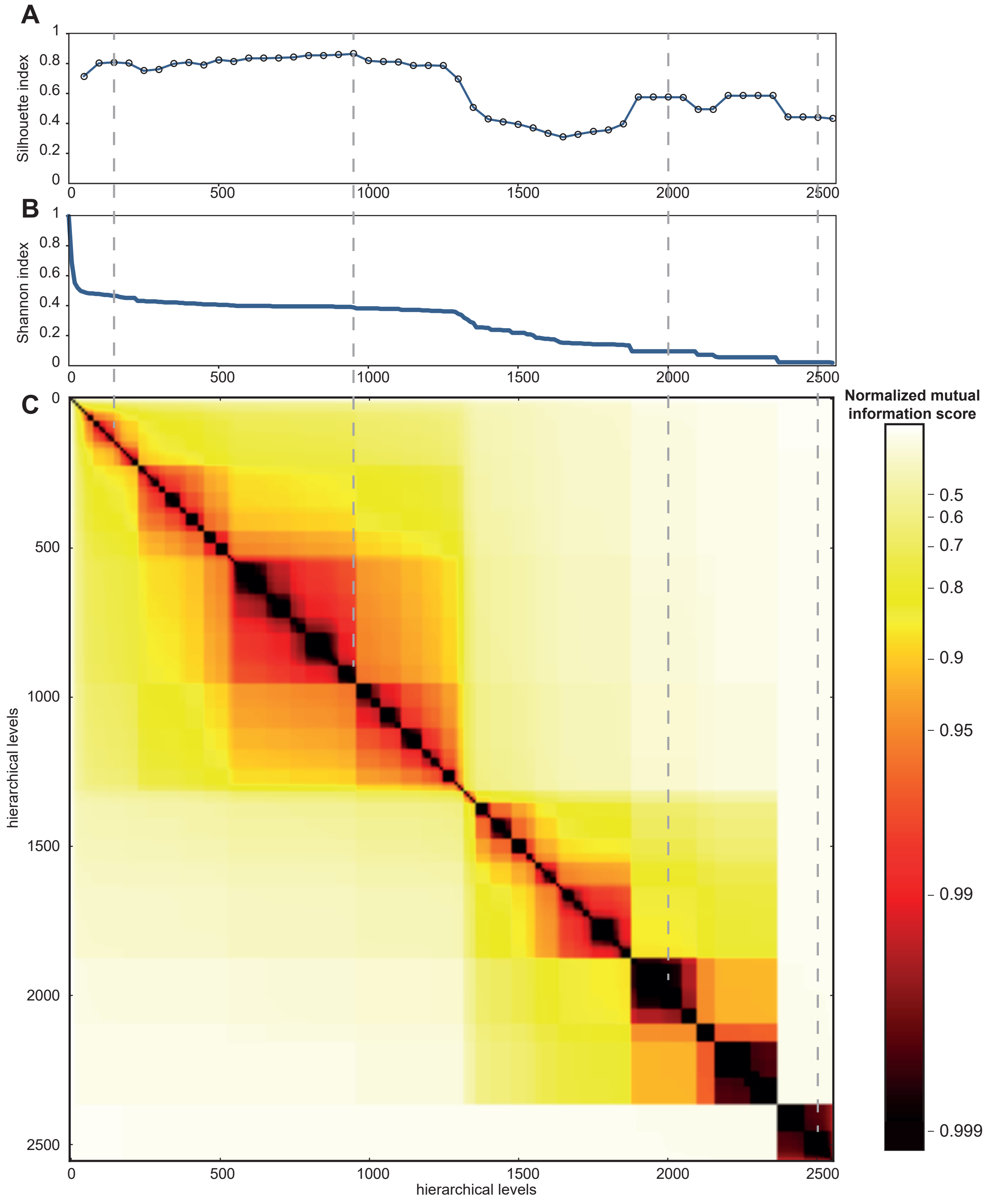


**Suppl. Figure 2**

### Supplementary Figure 3

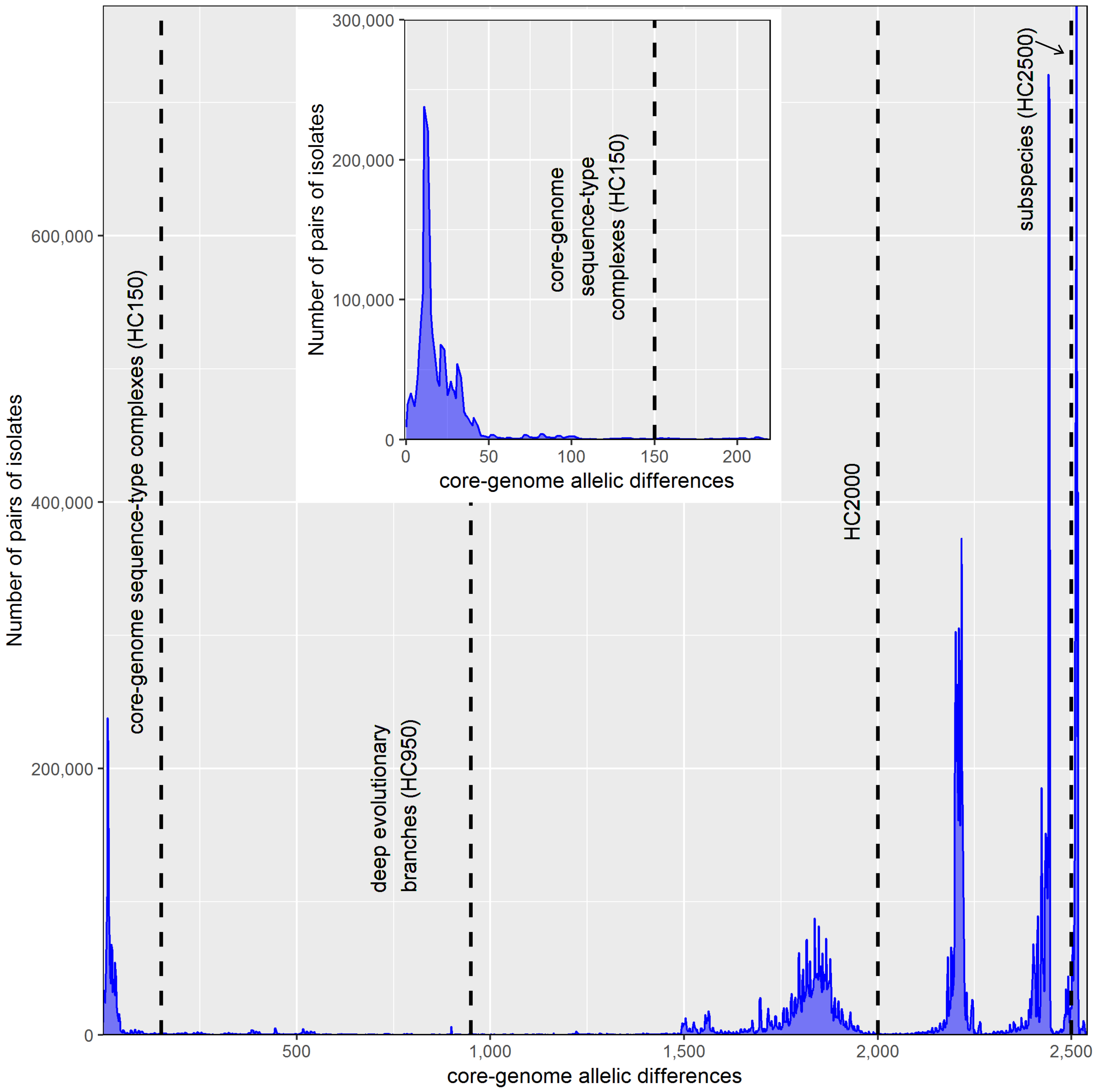


#### Suppl. Figure 3

### Supplementary Figure 4

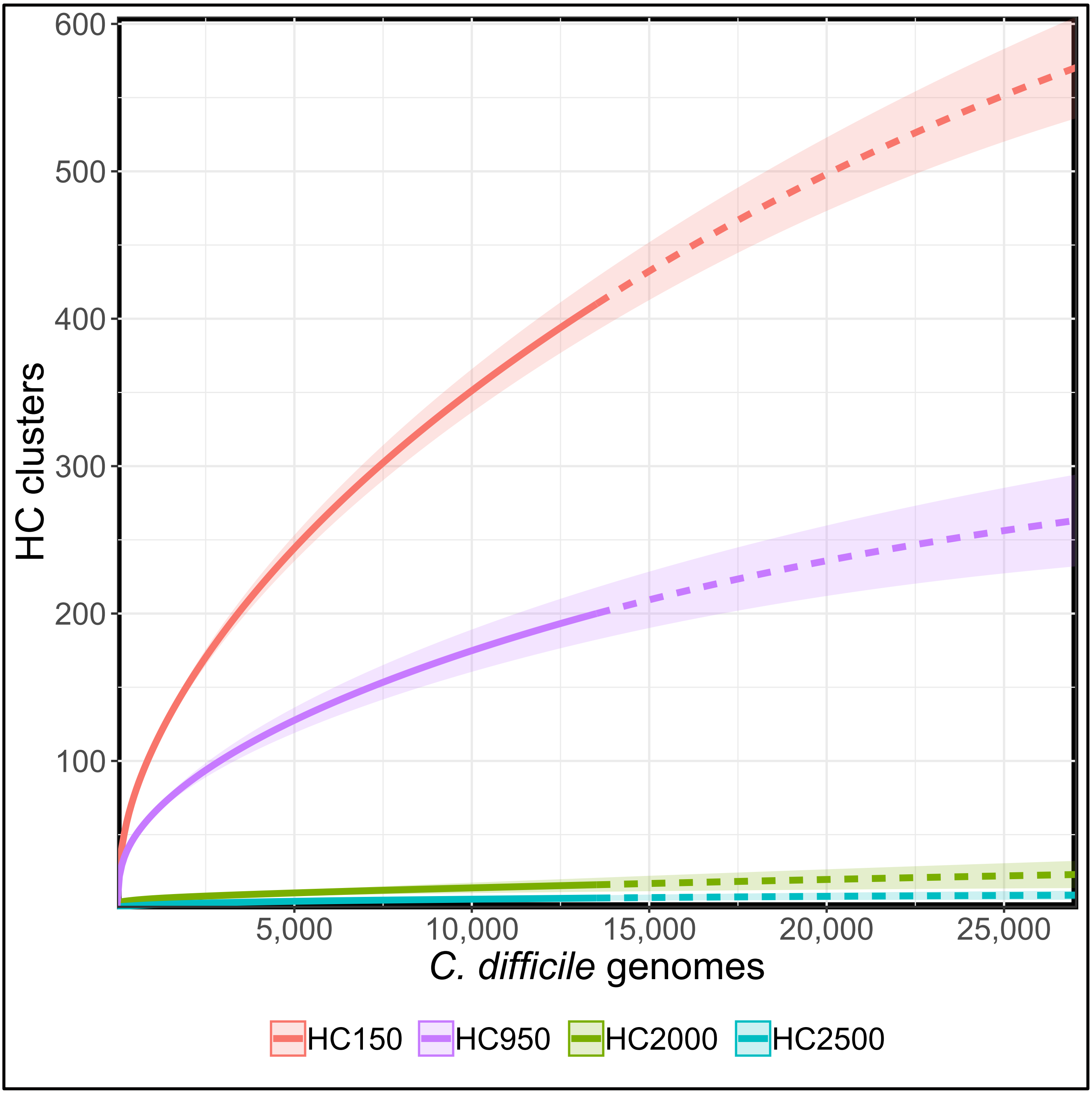


**Suppl. Figure 4**

### Supplementary Figure 5

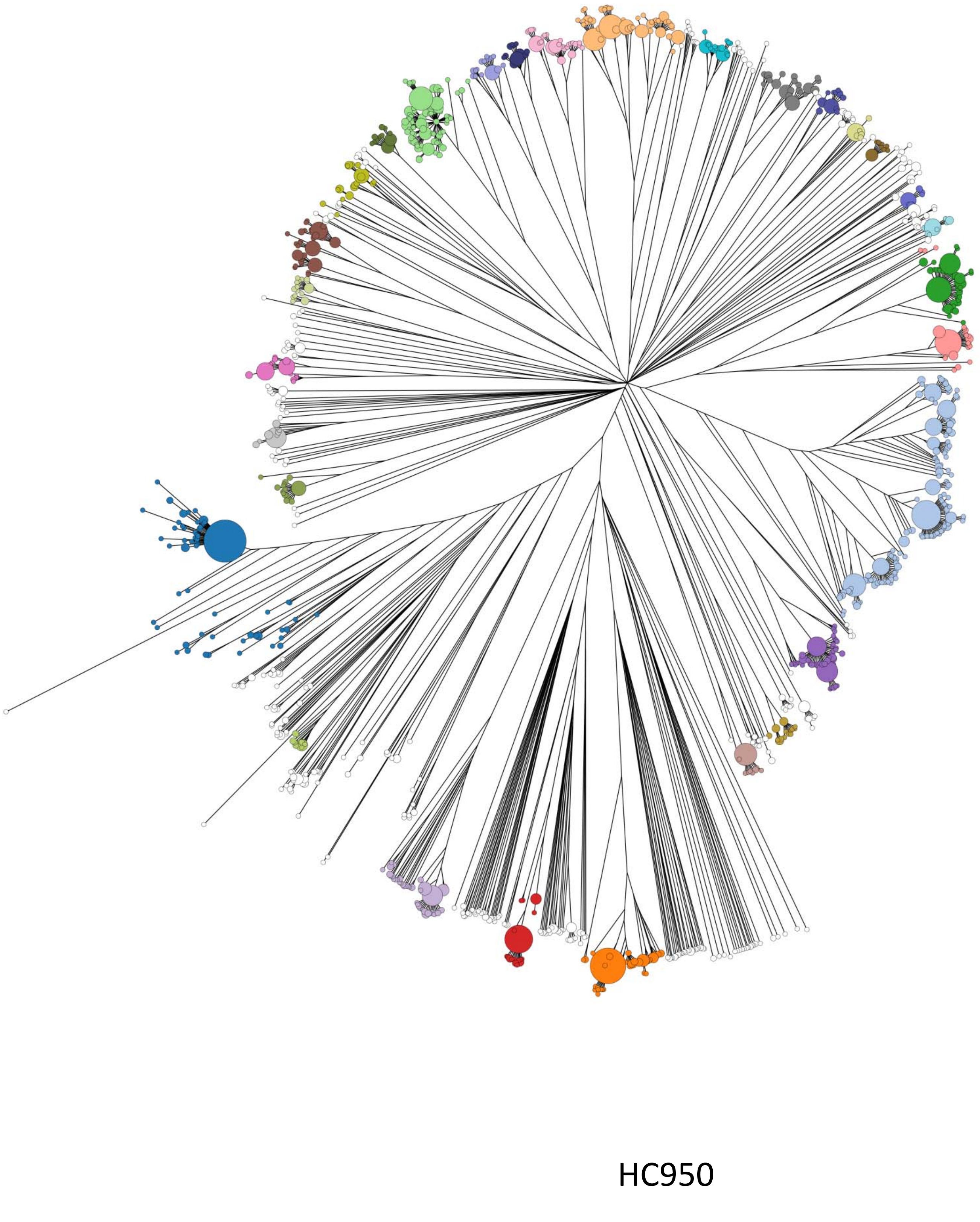


**Suppl. Figure 5**

### Supplementary Figure 6

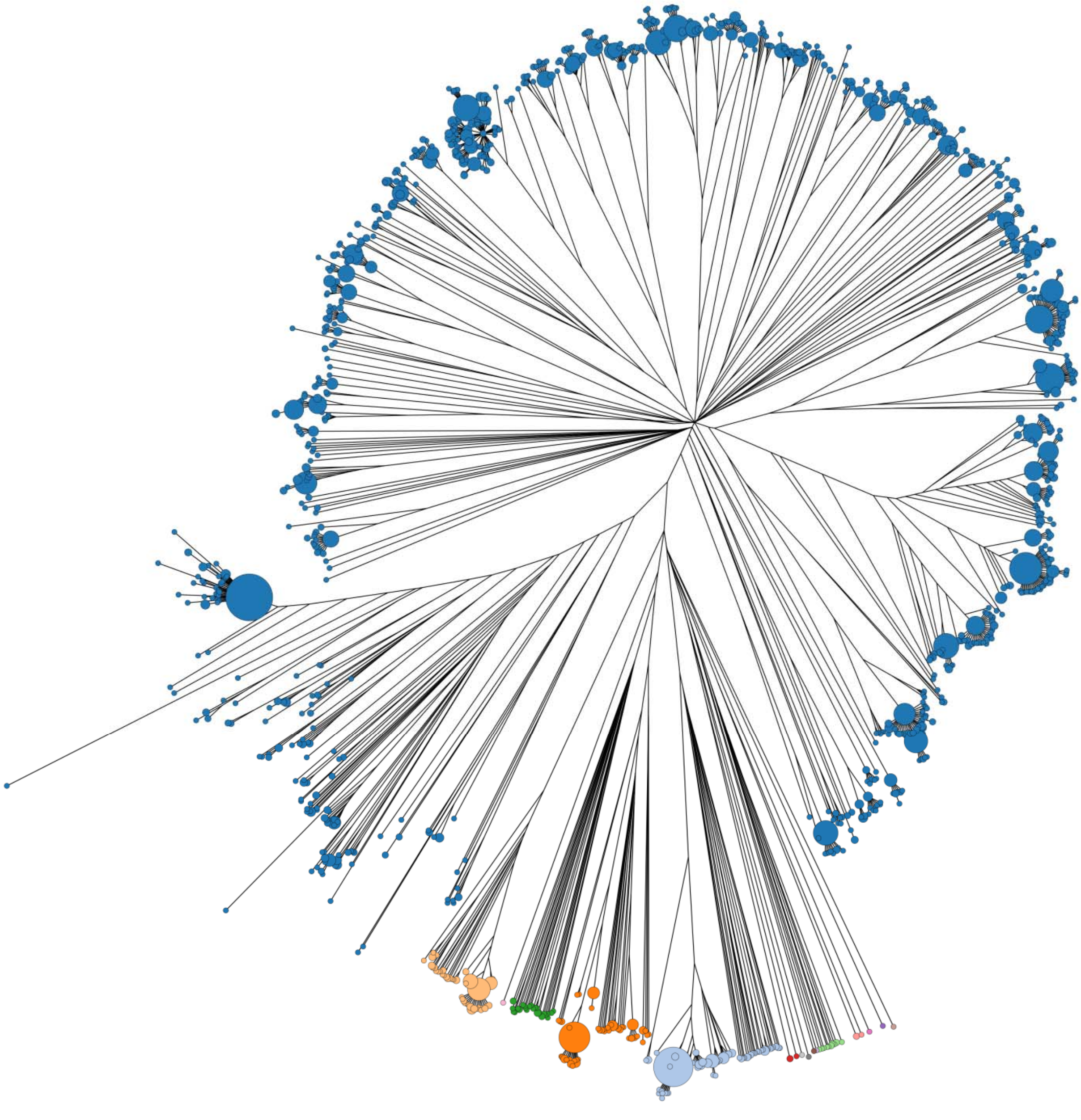

HC2000 ("super lineages")

**Suppl. Figure 6**

### Supplementary Figure 7

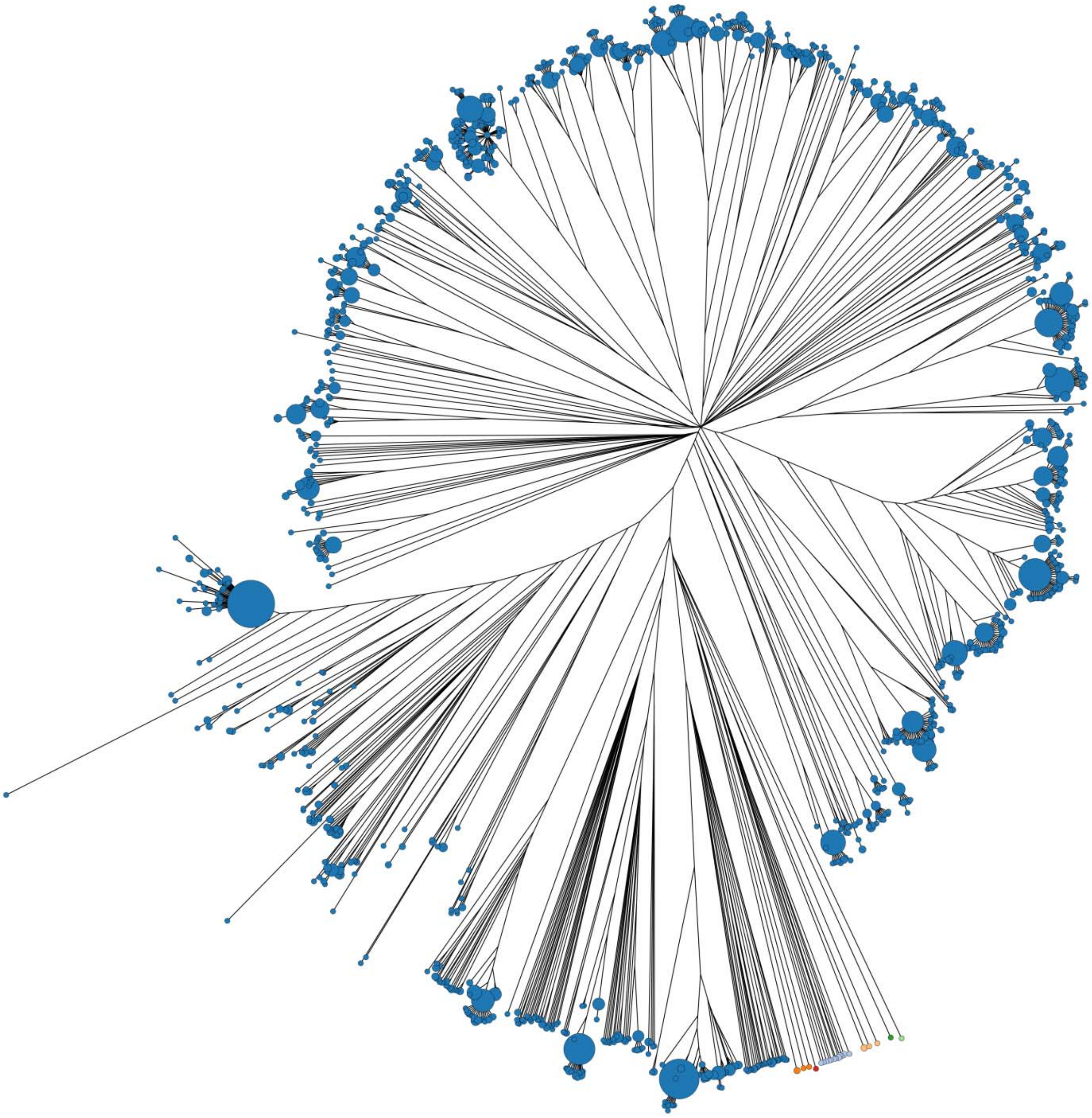

HC2500 ("subspecies")

**Suppl. Figure 7**

### Supplementary Figure 8

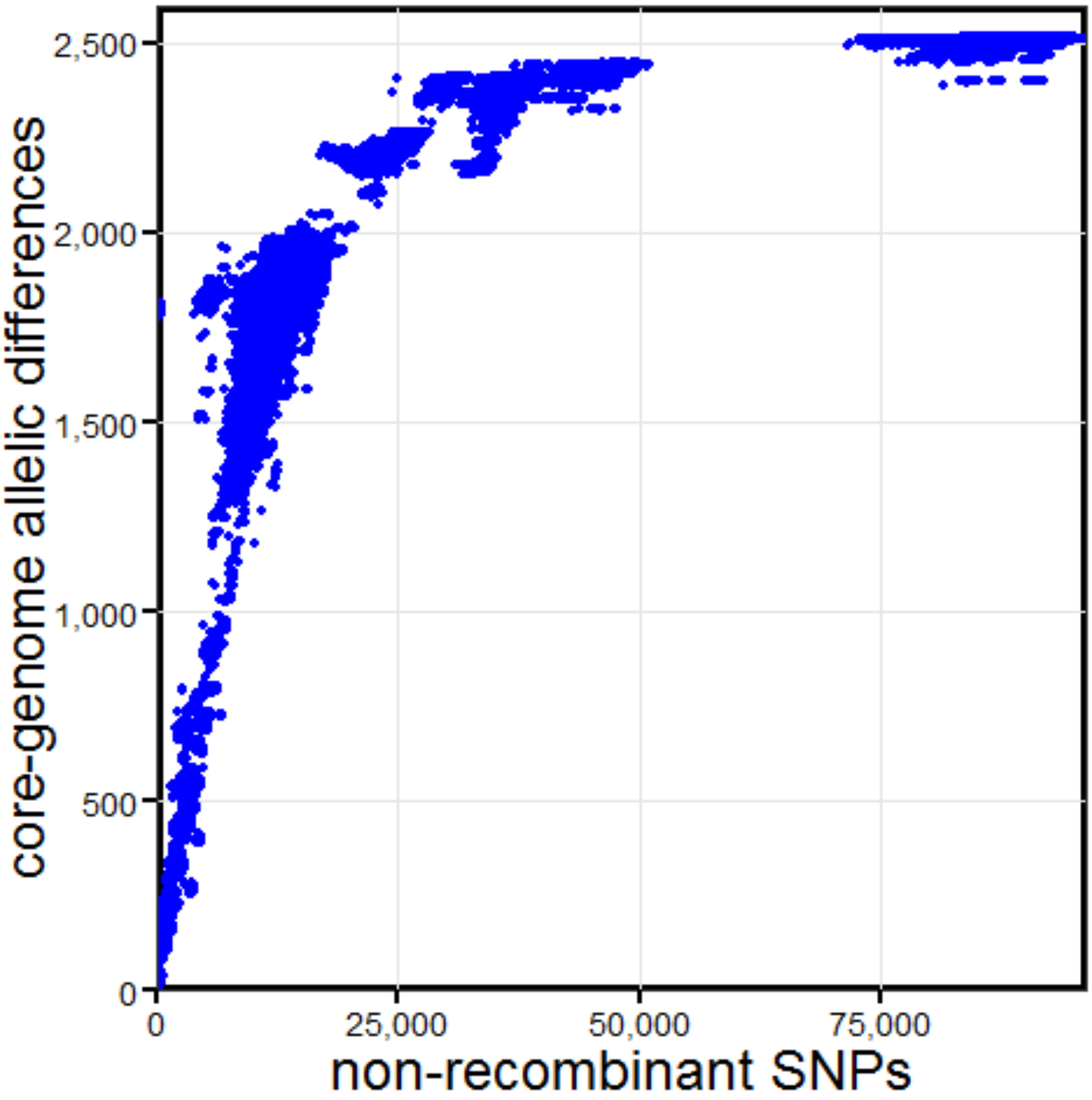


**Suppl. Figure 8**
